# Novel amylase genes enable utilisation of resistant starch by bifidobacteria relevant to early-life microbiome development

**DOI:** 10.1101/2024.10.09.617373

**Authors:** Molly E. Millar, Miriam Abele, Hannah C. Harris, Todor T. Koev, Andrea Telatin, Raymond Kiu, Douwe Van Sinderen, Yaroslav Z. Khimyak, Christina Ludwig, Lindsay J. Hall, Frederick J. Warren

## Abstract

*Bifidobacterium* species and strains are key members of the gut microbiota, appearing soon after birth and persisting into adulthood. Resistant starch is an important dietary substrate for adult-associated bifidobacteria, where its fermentation supports host health. However, little is known about how different starch structures interact with bifidobacteria across various ages and ecological niches. To address this, we carried out detailed growth kinetics screening of *Bifidobacterium* reference strains and unique isolates from breast-fed infants, testing their metabolic interaction with a variety of starch structures. ^1^H NMR metabolomics as well as analysis of CAZyme profiles from genomes were generated for each *Bifidobacterium*-starch combination. For a subset of resistant starch-utilising isolates, we integrated multi-omics approaches to attain further mechanistic interaction insights. Our results revealed that bifidobacterial starch hydrolysis capabilities are closely associated with their CAZyme profiles and appear to be connected to the niche they occupy. Notably, in one isolate of *Bifidobacterium pseudolongum*, we identified a novel gene cluster containing three multi-functional amylase enzymes complemented by several starch binding modules which were significantly upregulated in response to resistant starch. This gene cluster was also found in the genomes of bifidobacterial isolates from weaning infants and adults. These findings provide new insights into their participation in the maturation process of the infant gut microbiota. Uncovering mechanisms of metabolic interaction between starch structures and bifidobacteria underscores the importance of this ecological function and potential health implications.

## Introduction

The rapid taxonomic and functional changes in the human gut microbiota during early life have long-term impacts on health [1]. Gut microbiome assembly begins shortly after birth, and is influenced by a variety of factors such as gestational age, birth mode, exposure to probiotics or antibiotics, and infant diet [2]. Bifidobacteria are early colonisers of the infant gut, playing a key role in health [3–5]. For breastfed infants, human milk oligosaccharides (HMOs), which cannot be indigested by the host, are selectively fermented by certain *Bifidobacterium* species or strains, contributing to the infant’s microbiome development [2, 6, 7]. The weaning period, when solid foods are introduced, presents a critical stage for microbiome maturation [8]. This transition often involves starchy foods including cereals and vegetables [9] which contain resistant starch (RS) – a type of starch which is incompletely digested in the stomach and small intestine and reaches the colon where it can be fermented by gut microbes [10]. Since the intestinal microbiome of infants is underdeveloped, starch fermentation during weaning plays a role in gut microbiome ecological dynamics and maturation [9]. Fermentation of RS and other indigestible dietary components produces beneficial metabolites, such as short chain fatty acids (SCFAs) including acetate and butyrate [11], thereby contributing to various immunomodulatory effects which may continue into adulthood [12, 13]. This has led to harnessing the health-promoting attributes of bifidobacteria as probiotics for infants, particularly in preventing gastrointestinal conditions such as necrotising enterocolitis [14].

As important members of the developing gut microbiome, a range of *Bifidobacterium* species interact with RS as it moves through the digestive tract [15]. In both humans and several animal species, bifidobacteria can act as primary degraders of starch and facilitate cross-feeding with other gut microbes, influencing overall gut ecology [16–18]. Notably, an *in vitro* study using infant faecal inocula demonstrated an increased relative abundance of *Bifidobacterium* post-weaning in the presence of a high amylose maize starch, supportive of their ability to adapt to starch fermentation during weaning [19]. During this transition, *Bifidobacterium* must utilise at least one of the following: HMOs; solid food-derived glycans like starch, and/or host-derived mucins. *Bifidobacterium longum* (including subsp. *infantis* and subsp. *longum)* are particularly key to the transitional weaning period and there is emerging evidence of intra-species variation and nutrient-specific adaptation of bifidobacterial strains [15, 20, 21].

Starch granules consist of amylose and amylopectin, and their structural properties – such as crystalline structures [22] and amylose content (>30%) [23] – impacts their resistance to enzymatic attack and hydrolysis. Maize-derived starches with their varying amylose content including normal maize starch (NMS, 20% amylose), high amylose maize starch (HAM, 50% amylose), and HylonVII® (70% amylose) [24], offer a model for studying RS degradation. RS degradation requires multiple specialized enzymes and carbohydrate binding modules (CBMs) that target highly crystalline supramolecular structures, and break down a(1→4) and a(1→6) glycosidic linkages in amylose and amylopectin [18, 25–27]. While certain bifidobacterial species/strains, including those with the *apuB* gene encoding a bifunctional amylopullulanase, have been identified as starch degraders, the molecular mechanisms underlying RS utilisation remain understudied [28]. Identifying mechanisms of utilisation, and functional conservation in the genus, including species and strains from various ecological origins, could have implications for understanding the role of *Bifidobacterium* in the weaning gut microbiome.

To address these knowledge gaps, we investigated RS utilisation in eleven bifidobacterial strains/species from different ecological niches, including isolates from human infants and ruminants. We tested the general ability of these strains to grow on a range of structurally diverse starch substrates and performed detailed growth kinetics and metabolite release studies on a subset of six strains. We further explored the molecular basis of starch utilisation through genome annotation of hydrolytic enzymes and CBMs, completed by 3D protein and functional predictions. In two closely related animal isolates of *Bifidobacterium pseudolongum* we used transcriptomics and proteomics to interrogate gene regulation in response to RS. These experiments reveal how bifidobacteria-starch interactions could impact the gut microbiota, particularly during weaning, and uncover a novel gene cluster involved in RS degradation.

## Materials and Methods

### Bacterial strains

Strain accession numbers and source are described in Supplementary Table S1. Unique bifidobacteria (LH strains) were isolated as described previously [3]. A total of six bacterial strains were provided by the Hall Lab which were acquired as unique isolates from infant stool from the Baby-Associated MicroBiota of the Intestine (BAMBI) study. The associated metadata and research were published [29]. Notably, isolates tested in the present study were obtained at different ages of the same infant: *B. longum* subsp. *longum* LH277 at 37 days after birth; *B. pseudocatenulatum* LH656-662 at day 159 (all from the same stool sample, ANI values between strains >0.9999; Supplementary Table S3). *B. breve* LH24 was isolated from a different infant at 174 days after birth. Three infants (V1, V2, V3) donated at a similar age enabling isolation of 19 unique isolates. Reference strains used were purchased from suppliers NCIMB (Aberdeen, UK), UCC (Cork, Ireland), and ATCC (Manassas, VA, US).

### Starch growth assays

All strains were grown at 37 °C in either RCM, MRS media, or mMRS with specified carbohydrates (substituting glucose) in an anaerobic chamber (Don Whitley Scientific, Bingley, UK or Ruskinn Concept Plus, Bridgend, UK) containing 5% CO_2_,10% hydrogen, 85% nitrogen gas. All solid and liquid media used were supplemented with 0.05% (w/v) L-cysteine HCl and adjusted to pH 6.8 using concentrated NaOH. Media was sterilised to 126 °C for 15 min in a Prestige 2100 Classic Autoclave.

For starch media, three maize starches were selected with increasing amylose contents: unmodified regular corn starch (S4126; Sigma-Aldrich), practical grade high-amylose corn starch (S4180; Sigma-Aldrich, UK), HylonVII® maize starch (03451B00; gifted by Ingredion Inc, Manchester, UK). All starches were prepared by boiling starch suspensions at 100 °C in mMRS in a microwave at high power, mixing at regular intervals, for 1-3 min before being sterilised by autoclaving at 126 °C for 15 min. Experimental culturing involved first culturing in Reinforced Clostridial Medium (RCM) or MRS and strains were then sub-cultured using a 1:100 dilution in mMRS containing specified bacterial growth substrates. mMRS was used in the absence of any substrate as a negative control. To generate bacterial growth curves, 10 µl samples were removed for serial dilution at various time points in sterile MRS and plated on 90 mm petri dishes. The plates were incubated anaerobically in a Ruskinn Concept Plus for 2-3 days, and colonies counted. Data shown are mean values from three biological replicates.

### Bacterial metabolomics

Culture medium aliquots of 600 µl were stored in 1.5 ml snap-cap plastic microtubes at - 20 °C (if not immediately processed). If frozen, the aliquots were thawed at room temperature or in 4 °C overnight. Each sample was centrifuged at 13,000 rpm for 3 min and 400 µl of the supernatant was pipetted in NMR tubes (Norell® Standard Series™, 5 mm) followed by 200 µl NMR buffer, as previously described [30]. Quantification of metabolites was carried out on a Bruker Avance III NMR spectrometer, operating at a ^1^H frequency of 500.11 MHz equipped with an inverse triple resonance z-gradient probe. Spectra recorded were obtained using two different pulse sequences, used interchangeably due to data output being identical, Bruker’s ‘*noesygppr1d*’ and ‘*zggppewg’*. All other parameters were: ^1^H rr/2 *rf* pulse of 10 µs, a minimum of 128 scans, spectral width of 8000 Hz, acquisition time of 4 s, and relaxation delay of 10 s [30]. Spectra were apodized at 0.3 Hz, fitted and analysed using NMR Suite Processor v8.41 (Chenomx®, Edmonton, Canada).

### Proteomic data acquisition and data analysis

*B. pseudolongum* subsp. *globosum* NCIMB 702245 was cultivated for 24 h as described in 10 ml MRS and mMRS media. mMRS was supplemented with carbohydrates: 0.5% D-glucose (Sigma-Aldrich); cooked starch substrates prepared as described in mMRS with 1% normal maize starch (Sigma-Aldrich). Cultures were centrifuged for 10 min at 4,000 x *g*. The supernatant was discarded, 5 ml PBS was added to wash the cells. The samples were centrifuged for 10 min at 4,000 x *g* and supernatant was discarded. The cells were resuspended in 100 µL 100% TFA and transferred to clean plastic microtubes which were incubated in a thermocycler for 5 min at 55 °C. A volume of 900 µL 2 M Tris solution (pH was not adjusted) was added and tubes were vortexed. The lysed samples were quantified by Bradford assay and stored at −80 °C. Samples were processed for mass spectrometry according to the SPEED protocol as described previously [31, 32]. Briefly, a mass of 50 µg of protein per sample was reduced (10 mM TCEP) and carbamidomethylated (55 mM CAA) for 5 min at 95 °C. The proteins were digested with trypsin (proteomics grade, Roche) at a 1:50 enzyme:protein ratio (w/w) and incubation at 37 °C overnight. Digests were acidified with 3% (v/v) formic acid (FA) and desalted using self-packed StageTips (five disks per micro-column, 0 1.5 mm, C18 material, 3 M Empore) [33]. The peptide eluates were dried to completeness and stored at −80 °C. Each experiment was performed using three biological replicates.

Peptides were resuspended in 12 µl 0.1% FA in HPLC grade water and analysed on a Vanquish Neo coupled to an Orbitrap Exploris 480 mass spectrometer (both Thermo Fisher Scientific). Around ten µg of peptides were applied onto a commercially available Acclaim PepMap 100 C18 column (2 µm particle size, 1 mm ID × 150 mm, 100 Å pore size; Thermo Fisher Scientific) and separated using a stepped gradient with acetonitrile concentration ranging from 3% to 24% to 31% solvent B (0.1% FA, 3% DMSO in ACN) in solvent A (0.1% FA, 3% DMSO in HPLC grade water). A flow rate of 50 µl/min was applied. The mass spectrometer was operated in data-dependent acquisition (DDA) and positive ionization mode. MS1 full scans (360–1300 m/z) were acquired with a resolution of 60,000, a normalized automatic gain control target value of 100%, and a maximum injection time of 50 ms. Peptide precursor selection for fragmentation was carried out at a 1.2 seconds cycle time. Only precursors with charge states from two to six were selected, and dynamic exclusion of 30 s was enabled. Peptide fragmentation was performed using higher energy collision-induced dissociation and normalized collision energy of 28%. The precursor isolation window width of the quadrupole was set to 1.1 m/z. MS2 spectra were acquired with a resolution of 15,000, a fixed first mass of 100 m/z, a normalized automatic gain control target value of 100%, and a maximum injection time of 40 ms.

Peptide identification and quantification was performed with MaxQuant (version 1.6.3.4) with its built-in search engine Andromeda [34, 35]. MS2 spectra were searched against the NCBI *Bifidobacterium pseudolongum* subsp. *globosum* DSM20092 protein database (1882 protein entries, downloaded August 2022m protein duplicates were removes), supplemented with common contaminants (built-in option in MaxQuant). Trypsin/P was specified as proteolytic enzyme. Carbamidomethylated cysteine was set as fixed modification [36]. Methionine oxidation and acetylation at the protein N-terminus were specified as variable modifications. Results were adjusted to a 1% false discovery rate on peptide spectrum match (PSM) and protein levels. Label-Free Quantification (LFQ) and iBAQ intensities were computed [37].

### Alpha-amylase protein structure prediction

The full-length structure of multi-modular protein discovered in *B. pseudolongum* subsp. *globosum* NCIMB 702245 (*B. globosum* 45) containing alpha-amylase and CBM74 modules (designated as BpAmy74) was predicted using AlphaFold2 [1]. The colour-coded domain boundaries were determined using dbCAN [2] and InterProScan [3]. The BpAmy74 CBM74 domain was structurally aligned with the X-ray crystal structure of CBM74 from *Ruminococcus bromii* Sas6 with maltodecaose ligand (PDB ID: 7UWV), with RMSD = 1.039 Å (1325 to 1325 atoms), using UCSF ChimeraX software (version 1.5) [4]. The 7UWV structure was hidden except for the ligand. InterProScan was additionally used for classification of protein function and structure of multiple proteins in the gene cluster [38, 39].

### RNA extraction and sequencing

RNA extraction of *B. pseudolongum* subsp. *globosum* NCIMB 702245 were performed by culturing bacterial strains in 40 ml mMRS supplemented with: 0.5% D-glucose (Sigma-Aldrich), cooked starch substrates 1% w/v high amylose maize starch HylonVII, and 1% w/v normal maize starch.

A 2 ml sample was reserved for RNA extraction and transcriptome profiling of the bacterial cells in their initial transcriptional state (at 0 h) at point of inoculation. At 12 h and 24 h, parallel culture tubes were sacrificed for sampling 2 ml of the starch and glucose cultures, and 10 ml of the negative control. The samples were centrifuged at 4000 g for 10 min. The supernatant was discarded and 100 µl of RNAlater™ Stabilisation Solution (Invitrogen™, Thermo Fisher Scientific™) was used to resuspend the cell pellets. The suspensions were stored at 4 °C for a maximum of 14 h before RNA extraction. RNeasy® Mini kits (Qiagen, Manchester, UK) were used to perform the RNA extraction using the manufacturer’s protocol with the following additional chemical lysis and bead beating steps. A volume of 10 µL β-mercaptoethanol (Sigma-Aldrich) was added to 1 ml Buffer RLT Plus before use. Bacterial cells containing RNAlater™ (Thermo Fisher Scientific™) were mixed with 600 µL RLT lysis buffer and transferred to lysing matrix E-tube and placed in an MP Biomedicals™ Fastprep 24. Speed setting 6 for 60 seconds was repeated 3 times (3 min total) keeping samples on ice between steps. Tubes were centrifuged for 10 min at 14000 g. The supernatant was transferred to a 2 ml Low-bind Eppendorf tube where 600 µL of 70% ethanol was added and mixed well by pipetting gently. A volume of 700 µL was then transferred to an RNeasy spin column and the protocol was continued using the manufacturer’s protocol. A volume of 35 µL RNase free water was added to the spin column membrane to elute the RNA. Samples were incubated for 1 min at room temperature before being centrifuged at 8000 x *g* for 1 min – the spin column was discarded after eluate was obtained. TURBO DNA-free™ kit for DNA removal was used following the manufacturer’s protocol.

Purified RNA was quantified and quality controlled using an Agilent 4200 TapeStation using High Sensitivity RNA ScreenTape according to the manufacturer’s protocol. All samples were ensured to have RIN^e^ values above 8.0. RNA sequencing was performed at Azenta, Cambridge, UK, where rRNA depletion was performed using the NEBNext rRNA Depletion Kit (Bacteria). RNA sequencing library preparation was performed using NEBNext Ultra II RNA Library Prep Kit for Illumina by following the manufacturer’s recommendations (NEB, Ipswich, MA, USA). Briefly, enriched RNAs were fragmented according to manufacturer’s instruction. First strand and second strand cDNA were subsequently synthesized. cDNA fragments were end repaired and adenylated at 3’ends, and universal adapter was ligated to cDNA fragments, followed by index addition and library enrichment with limited cycle PCR. Sequencing libraries were validated using NGS Kit on the Agilent 5300 Fragment Analyzer (Agilent Technologies, Palo Alto, CA, USA), and quantified by using Qubit 4.0 Fluorometer (Invitrogen, Carlsbad, CA). The sequencing libraries were multiplexed and loaded on the flow cell on the Illumina NovaSeq 6000 instrument according to manufacturer’s instructions. The sequencing parameters selected were depth of ∼20 million paired end, 150bp reads per sample. The samples were sequenced using a 2×150 Pair-End (PE) configuration v1.5.

Image analysis and base calling were conducted by the NovaSeq Control Software v1.7 on the NovaSeq instrument. Raw sequence data (.bcl files) generated from Illumina NovaSeq was converted into fastq files and de-multiplexed using Illumina bcl2fastq program version 2.20. One mismatch was allowed for index sequence identification.

### Transcriptome data processing

The raw reads were delivered in FASTQ format and were quality checked with SeqFu v1.19.0 [40], discarding reads with undetermined bases, or shorter than 100 bp, and filtered with SortMeRNA v4.3.6 [41]. The samples were processed by read mapping with a reference genome (GCF_002706665.1_ASM270666v1_genomic.fna) using RNAFlow [42] v1.4.1 executed on an HPC with Slurm by Nextflow v21.10.6 [43, 44], using BamToCov [45] to support prokaryotic GFF annotations and skipping SortMeRNA as it was done previously.

The pipeline performed a second QC using fastp v0.20.0 [46] then mapped the reads against the reference genome using HISAT2 v2.1.0 [47]. Read mapping against each of the coding sequences (CDS) was counted with BamToCov v2.7, producing a raw counts matrix which was then analysed with DESeq2 [48] to identify differentially expressed features.

### Statistical analyses

Statistical analysis of bacterial growth data was carried out using area under the curve calculations and repeated measures 2-way ANOVA comparing starch to the negative control at each time point, performed in GraphPad Prism v9 (GraphPad Software). Protein-Groups.txt result file from MaxQuant was used as an input for Omicsviewer 1.1.5 [49] to perform t-tests of differential protein abundances of *B. pseudolongum* subsp. *globosum* NCIMB 702245 in the presence of normal maize starch or glucose. For RNA expression analysis, using iDep.96, data were visualised and DESeq2 was used to generate differential expression values for each gene and associated FDR values [48, 50]. All other statistical analyses were performed in GraphPad Prism v9.

### Whole bacterial genome sequencing

For genome sequencing of a reference type strain purchased, *B. pseudolongum subsp. pseudolongum* DSM 20099 / NCIMB 702244, the strain was cultured from 30% v/v glycerol stock. The strain was sub-cultured using 500 µl in 10 ml in MRS and incubated anaerobically for 48 h at 37 °C in a Don Whitely Scientific miniMACS anaerobic incubator for each incubation. The culture was centrifuged for 10 min at 4000 x *g*. The supernatant was discarded, and 5 ml PBS was added to resuspend the bacterial pellet. 200 µl of this bacterial solution was added to an MP Biomedicals™ Lysing Matrix E tube and the protocol was followed as previously described [51]. The method utilises FastDNA™ SPIN Kit for Soil, following manufacturer instructions, with extended 3 min bead-beating. The DNA extracts were stored at −20 °C.

For DNA sequencing using Oxford Nanopore MinION, primers were designed with a 3’ end compatible with the Nextera transposon insert and a 24bp barcode at the 5’ end including a pad and spacer. The same barcode was used at each end using 96 combinations. Primer pairs were mixed at a concentration of 10 µM. The tagmentation reaction was reduced 20-fold. A Tagmentation Mix was prepared by combining three reagents: TB1 Tagment DNA Buffer (0.5 µl) was mixed with 0.5 µl bead-linked transposomes (BLT) (Illumina) and 4 µl PCR grade water. The total volume prepared (5 µl) was added to a chilled 96-well plate. DNA was normalised to between 50-100 ng/µl. Thereafter, 2 µl of normalised DNA was pipette mixed with the 5 µl of the Tagmentation Mix and heated to 55 °C for 15 minutes in a PCR thermoblock.

A PCR master mix was made using a ratio of 10 µl NEB LongAmp Taq 2X Master Mix to 2 µl PCR grade water (12 µl required per sample). To each well of a new 96-well plate, 12 µl of the master mix was added. Additionally, 1 µl of the appropriate primer pair at 10 µM was added to each well. Finally, the 7 µl of Tagmentation mix was added and mixed by pipette.

The PCR was run using the following parameters: 1 cycle at 94 °C for 2 min; 14 cycles at 94°C for 30 sec; 55 °C for 1 min; 65 °C for 15 min; 1 cycle at 65 °C for 15 min; and finally hold set at 10°C. Following the PCR reaction, the libraries were quantified according to the manufacturer’s instructions using QuantiFluor® dsDNA System (Promega) and run on a GloMax® Discover Microplate Reader (Promega). Libraries were pooled together in equal nanograms and SPRI size selected at 0.6X bead volume using Illumina Sample purification beads with final elution in 32 µl. A 0.75% Agarose Cassette was used to run 30 µl of sample for processing using a sageELF (Cat. No. ELF0001, Sage Science, Beverly, USA). sageELF fractionates the DNA using electrophoresis to separate the fragments based on size. Using a running time of 3 hours and 40 minutes on the 1-18kb run mode, 10kb fragments were in the middle of the 12 elution wells. The eluted fractions were then run on an Agilent 4200 TapeStation using D5000 Genomic ScreenTape (Agilent); fragments of 3kb and above were pooled together. Pooled fractions underwent a 1X SPRI clean-up and were eluted in 50 µl.

The eluate then underwent genomic ligation protocol using a Ligation Sequencing Kit (Oxford Nanopore Technologies) and was loaded on a MinION device according to the manufacturer’s instructions for sequencing.

### Whole bacterial genome assembly

Using long reads from MinION sequencing data, a draft genome of a single contig containing 1,916,804 bp was generated as follows. Firstly, Filtlong v0.2.1 (installed from the BioConda channel [52]) was used to filter long reads by preferentially selecting higher quality, longer reads from the read set, in which the worst 10% reads and reads <1kbp were discarded [53]. Next, de novo assembler Flye v2.9 (--nano-hq mode) was used to assemble the Nanopore long reads into a draft genome with 5 polishing iterations (with option -i 5) [54].

Subsequently, the genome assembly was further corrected and polished with Medaka v1.4.3 (Oxford Nanopore Technologies, Oxford, UK). A final tool, Circlator v1.5.5 was used attempting to circularise the genome assembly [55]. However, the final genome of one contig was not fully circularised. The WGS of *B. pseudolongum* 44 and *B. globosum* 45 were assessed for strain similarity (average nucleotide identity (ANI) score) using pyani v0.2.7 [56].

### Carbohydrate active enzyme and binding module genome annotation

Bacterial isolate whole genome sequences were obtained from the NCBI database [57] or sequenced in-house: the full isolate details and accession numbers are available in Supplementary Table S1; whole genome sequencing was performed for *B. pseudolongum* subsp. *pseudolongum* NCIMB 702244 according to described in previous section. For whole genome analysis of strains, dbCAN was used as a tool for prediction of glycan substrates for CAZymes, associated CBMs, and gene clusters (CGCs) by searching against the Cazy database [58].

## Results

### Bifidobacterium CAZyme profiles

*Bifidobacterium* strains tested using multiple starch structures were isolated from a range of ecological niches ranging from the preterm infant gut to the bovine rumen (Figure 1).

**Figure 1.**
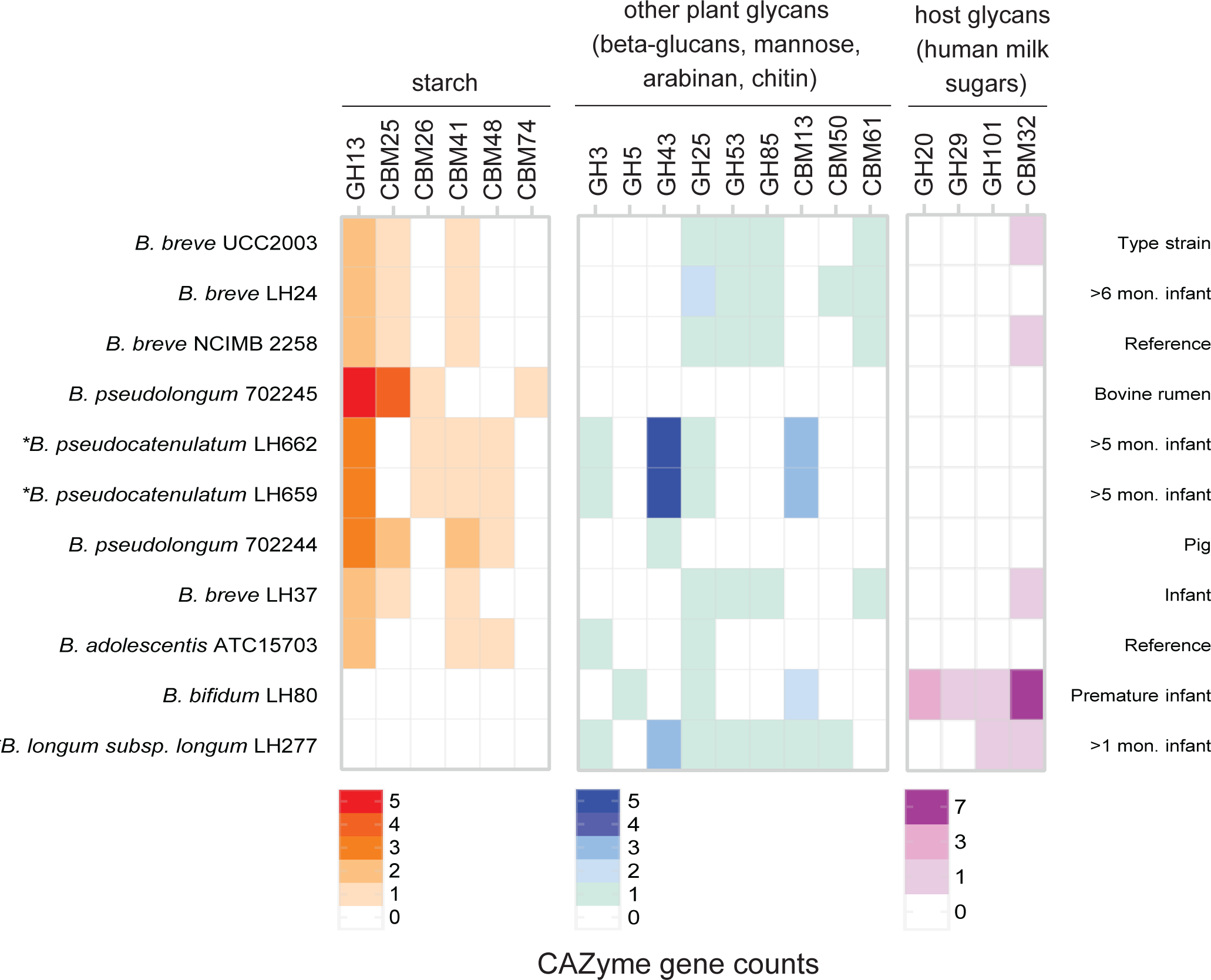
CAZyme profiles of *Bifidobacterium* isolates from different sources. The number of each CAZyme families and carbohydrate binding modules detected in the whole genomes of each strain was assessed and categorised by substrate (starch, other plant glycans, host glycans). Results shown are only CAZyme sequences detected which also contained a signal peptide implying that the enzymes would be extracellularly expressed. Asterisks denote that isolates were derived from stool donated from the same infant.

Glycosyl hydrolase (GH) families with enzymes involved in starch degradation includes GH 3, 13, 14, 15, 31, 57, 119, and 126 [59], of which the assessed isolates only encoded GH13 and GH3 family enzymes, the latter being more frequently associated with other plant glycan substrates (full CAZyme profiles can be found in Supplementary Table S2). Additionally, several starch-specific CBMs were identified including CBM 25, 26, 41, 48, and 74 in these strains. To focus on enzymes involved in extracellular starch hydrolysis, we filtered results to identify CAZymes with a signal peptide (Figure 1). Strains that lacked genes encoding a signal peptide-containing GH13 enzyme as well as relevant starch-related CBMs were isolated from very young (<1 month old) or preterm infants (*B. bifidum* and *B. longum* subsp. *longum*). Despite lacking starch-degrading capabilities, these isolates did possess sequences related to host glycan metabolism, particularly HMO degradation (Figure 1).

Amongst the remaining nine bacterial isolates, *B. pseudocatenulatum* from a weaning-age infant (5-6 months), and two animal isolates of *B. pseudolongum* were predicted to genomically encode the highest number of GH13 enzymes (Figure 1). The *B. pseudocatenulatum* isolate was also shown to encode several families involved in degradation of other plant glycans, but none related to host glycan metabolism. This suggests a shift in substrate utilisation as the gut microbiota adapts to dietary changes during weaning.

### Starch degradation by *Bifidobacterium* isolates from diverse ecological origins

*B. pseudocatenulatum*, *B. pseudolongum*, and *B. breve* strains were all able to hydrolyse multiple types of starch (Figure 2A), consistent with their CAZyme profiles. In contrast, 3 strains – *B. adolescentis* ATC15703, *B. bifidum* LH80, and *B. longum* subsp*. longum* LH277 – showed no significant growth on starch compared to control conditions (Figure 2A). We report an abnormal result for this *B. adolescentis* strain which is most commonly known as a starch-degrading species, especially since this strain encodes two extracellular GH13 family enzymes [18]. Amongst the various starch types tested, growth values varied, with HylonVII (a 70% amylose maize starch) being the least accessible for all strains except *B. breve* UCC2003. Only *B. pseudolongum* subsp*. globosum* NCIMB 702245 (henceforth referred to as *B. globosum* 45) was able to use normal maize starch (NMS) and HylonVII starch equally well. The deduced proteome of this strain (11 hits), alongside *B. pseudocatenulatum* isolates, encompasses a higher prevalence of GH13 hits and starch-binding CBMs compared to 4 hits for *B. breve* UCC2003. *B. globosum* 45 also encodes the RS-binding CBM74 [60].

**Figure 2.**
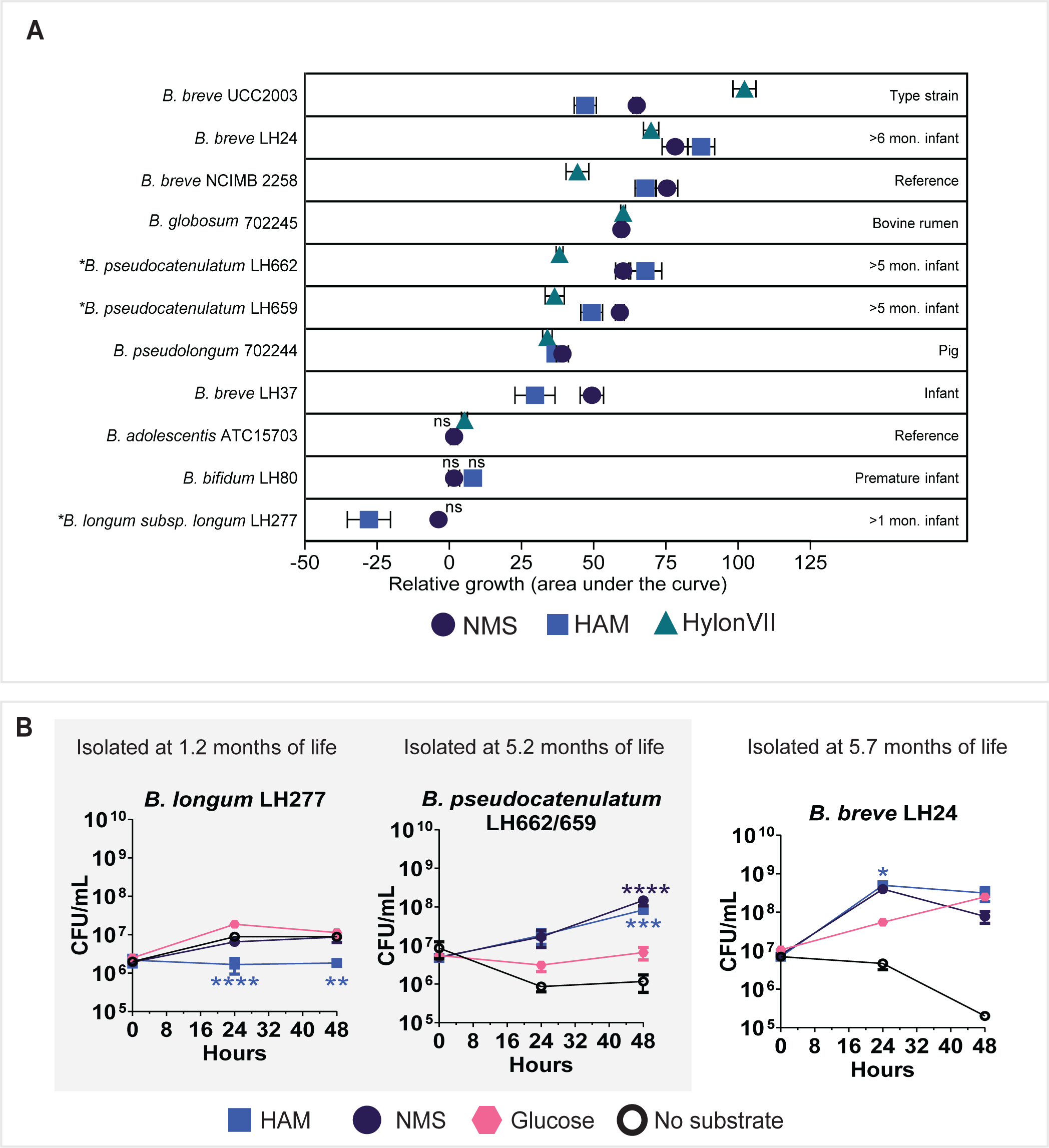
Growth of *Bifidobacterium* strains in the presence of different starches. (A) The growth of bacteria in the presence of starch was compared to growth in the absence of a substrate, where the mean difference (mean diff.) in the area under the curves (AUC) was statistically compared to assess starch utilisation in mMRS medium supplemented with 1% w/v: normal maize starch (NMS), high (50%) amylose maize starch (HAM), or HylonVII® (70% amylose) starch. The growth of each strain was determined by generating by log10 conversion of the CFU counts from 3-5 time points between 0-48 h and calculating the AUC (experiments were carried out in triplicate). The AUC and SEM were calculated and used to perform a Dunnett’s multiple comparisons test between the starch condition and the no substrate control. A positive integer value for mean difference between AUC relative to the negative control indicates growth on the starch substrate; a negative integer indicates no growth. Values are statistically significant unless indicated (ns >0.01). (B) Growth of *Bifidobacterium* isolates in the presence of 1% w/v normal maize starch (NMS), high (50%) amylose maize (HAM), 1% w/v D-glucose, and no substrate in modified MRS media. The isolates tested were obtained at different ages of the same infant: *B. longum* subsp. *longum* LH277 at 37 days after birth; *B. pseudocatenulatum* LH656-662 at day 159 (ANI values between strains >0.9999). *B. breve* LH24 was isolated from a different infant at 174 days after birth. All data were generated in triplicate. Dunnett’s multiple comparisons tests were performed to compare the CFU/ml mean values of the starch conditions to the no substrate (negative) control mean values. Significance values: * <0.05, ** <0.01, ** < 0.001, and **** <0.0001.

To explore the relationship between starch utilisation by a given bifidobacterial strain and corresponding host age, we compared growth kinetics of *B. pseudocatenulatum* LH662/LH659 and *B. infantis* LH277 isolated from the same infant at different time points (Figure 2B). *B. infantis* LH277 had significantly lower growth in the presence of high amylose maize (HAM) starch and no significant growth in the presence of NMS compared to the negative control with no added carbon source (Figure 2B). *B. pseudocatenulatum* and *B. breve* LH isolates, derived from infants’ stool samples collected during the weaning phase (∼5 months), were capable of degrading both NMS and HAM. Their growth was significantly higher compared to the control on at least one starch substrate, indicating an age-dependent adaptation to starch utilisation during weaning.

### Starch structure impacts growth rate and metabolite output of *Bifidobacterium* in a strain-dependent manner

The impact of starch structure on growth kinetics of *Bifidobacterium* isolates was analysed in combination with metabolomics. In the presence of NMS and HylonVII, *B. breve* UCC2003 was shown to exhibit significantly increased growth, ∼4X higher than the negative control (p <0.0001) (Figure 3, top left) (full statistical data available in Supplementary Table S3). By 48 h, cell counts in the presence of NMS were no longer significantly different to the control, whereas cells grown in HylonVII entered stationary phase, with counts still significantly higher than the control (p = 0.0115). The difference between cell counts between NMS and HylonVII was significant (p = 0.0117) at 48 hours suggesting that starch structure and amylose content significantly affected growth patterns. Maltose concentrations during HylonVII fermentation were significantly lower than during NMS fermentation (p = 0.0175). The acetate levels in the NMS condition were significantly higher by an order of magnitude compared to the negative control (p < 0.0001) and the glucose control (p = 0.0007), peaking at 18 hours. In contrast, acetate concentrations were significantly lower in the HylonVII condition compared to NMS at every time point after 18 h (p < 0.0001). Similar growth, acetate, and maltose utilisation patterns were observed in *B. breve* NCFB 2258 (Figure 3, top right), suggesting consistent starch utilisation phenotypes between these strains.

**Figure 3.**
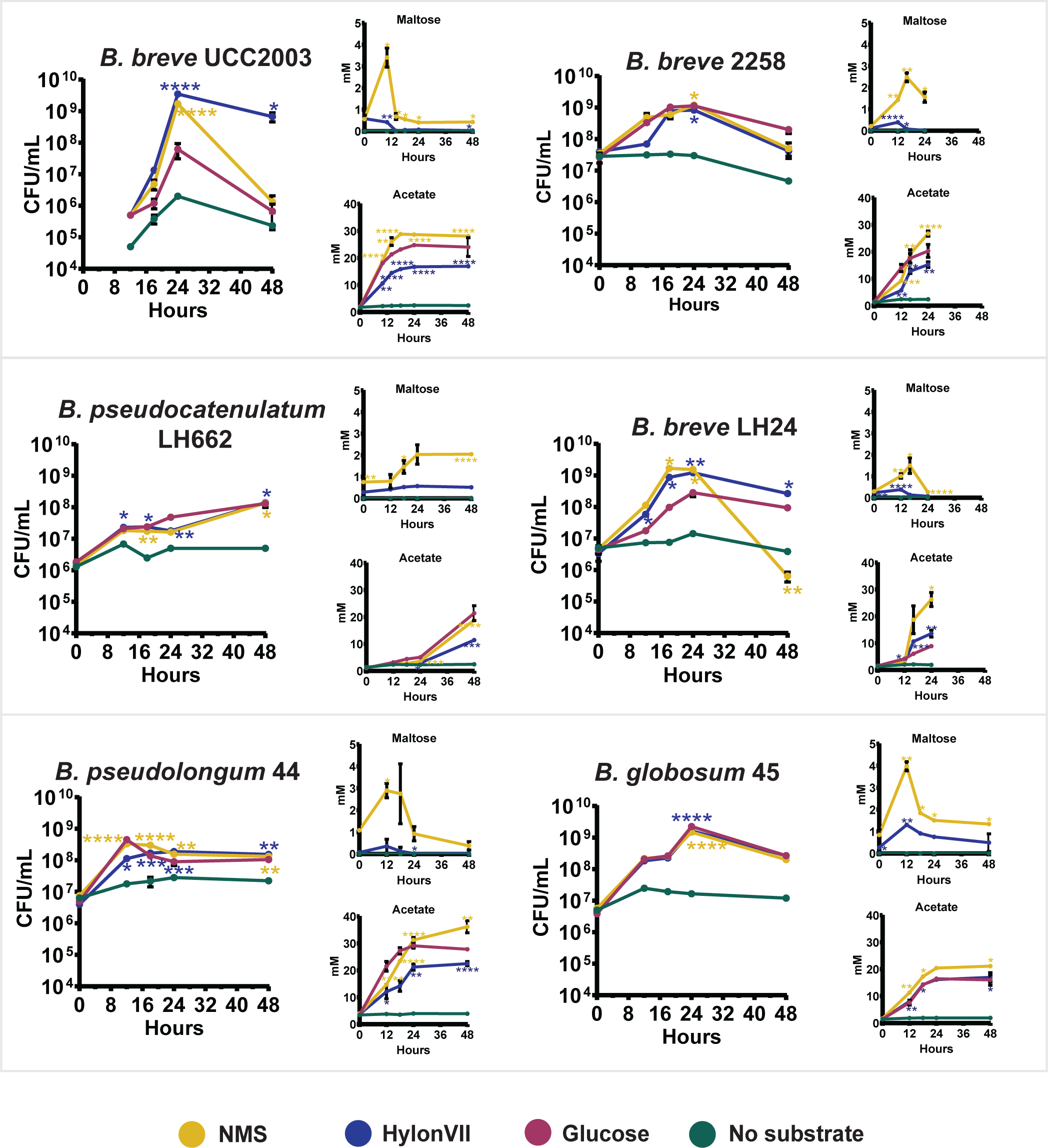
Variation in growth kinetics and metabolite production by Bifidobacterium strains. Acetate and maltose concentrations were quantified in the presence of no substrate (negative control), glucose, normal maize starch (NMS) and 70% amylose maize (HylonVII) starch. Asterisks represent a significant difference (using repeated measures ANOVA) between substrate at each individual time point where **** p<0.0001 ***p<0.001 **p<0.01 *p<0.05. Statistical models were used to ascertain if starch had a significant effect on the growth of each strain relative to the negative control. The same statistical methods were used to analyse the impact of each starch on production of maltose and acetate. Values displayed for each time point are the mean of three biological replicates.

In contrast, two unique isolates of *B. pseudocatenulatum* LH662 and *B. breve* LH24 derived from the stool of infants aged 5-6 months, exhibited different starch utilisation profiles. *B. pseudocatenulatum* LH662 showed significantly increased growth on each type of starch compared to the negative control, yet elicited a lower rate of increase compared to *B. breve* LH24 (Figure 3, middle left and middle right). This strain was in lag phase for the first 24 h followed by a gradual increase in growth towards 48 h. However, maltose concentration in the supernatant increased after 12 h and remained elevated over 48 h, suggesting partial starch hydrolysis without complete maltose uptake. In contrast, *B. breve* LH24 was more efficient in degrading resistant HylonVII, increasing its cell count by three orders of magnitude within 24 h. In the NMS condition, maltose peaked then rapidly declined by 24 hrs, leading to a higher acetate production to the HylonVII condition.

For the animal isolates, porcine isolate *B. pseudolongum* subsp. *pseudolongum* NCIMB 702244 (henceforth referred to as *B. pseudolongum* 44) and bovine rumen isolate *B. globosum* 45, we observed different utilisation phenotypes (Figure 3, bottom left and bottom right). *B. pseudolongum* 44 was shown to exhibit less growth on more resistant starch HylonVII compared to NMS (p < 0.0001 at 12 h (peak growth) time point), and this was reflected in its acetate production in the HylonVII condition (p = 0.0128 at end point of 48 h). In contrast, *B. globosum* 45 was able to utilise both starch types equally, with no significant difference in cell counts at 24 h (p = 0.7562 at 24 h (peak growth) time point), nor acetate concentrations (p = 0.1201). Additionally, *B. globosum* 45 exhibited two exponential growth phases, separated by a short lag phase, and released and consumed more maltose from HylonVII compared to *B. pseudolongum* 44. Whilst *B. globosum* 45 was able to more completely hydrolyse HylonVII starch, *B. pseudolongum* 44 achieved its peak growth at 12 h compared to 24 h for *B. globosum* 45. Moreover, lower end-point acetate concentrations in *B. globosum* 45 (36 mM vs 21 mM) suggest that, despite enhanced hydrolytic capacities, its efficiency in converting sugars into short-chain fatty acids was lower.

The variation in HylonVII utilisation between the two bifidobacterial isolates *B. pseudolongum* 44 and *B. globosum* 45 prompted an investigation into the genomic drivers behind their variation in growth kinetics and metabolite output. WGS analysis of *B. pseudolongum* 44 and *B. globosum* 45 indicated an average nucleotide identity (ANI) score of 93.4%, indicating that the strains are not members of the same species (nor subspecies), but are closely related [61]. This close relationship, combined with their phenotypic variation, presented an opportunity for further investigation into the manner by which they metabolize starch.

### Exploring genomic drivers of *Bifidobacterium*-starch interactions

To investigate the mechanisms underlying RS utilisation in *Bifidobacterium*, we then explored the genomic differences between *B. pseudolongum* 44 and *B. globosum* 45 to make predictions about the genes involved in their starch-degrading phenotype. WGS annotation using dbCAN revealed that *B. globosum* 45 had a higher number of GH13 family enzyme hits and starch-specific CBMs compared to *B. pseudolongum* 44 (Supplementary Table S5). Notable, *B. globosum* 45 possessed various multi-modular proteins with multiple CBMs associated with a single GH13 enzyme, many of which were predicted to be extracellular based on the presence of signal peptides (Figure 4A) Among the identified proteins, *B. globosum* 45 contained five predicted extracellular a-amylases, compared to just two for *B. pseudolongum* 44 (Supplementary Table S5). These a-amylases in *B. globosum* 45 included auxiliary starch binding modules such as CBM26 and CBM74 (WP_099309720.1, here referred to as alpha-amylase_720), which are absent in *B. pseudolongum* 44. One particular a-amylase exhibited 77.73% sequence identity in *B. pseudolongum* 44 indicating significant protein homology between the two subspecies; the main difference between the two proteins was absence of CBM74 module in *B. pseudolongum* 44. Interestingly, three of these multi-modular proteins were located on the same genomic locus (Figure 4A), and are not currently designated as a-amylases, but as ‘Ig-like domain containing proteins’ in the NCBI database (Supplementary Table S6). These three proteins (alpha-amylase_162, alpha-amylase_720, and alpha-amylase_110) are clustered at the same locus, with alpha-amylase_720 as the largest central protein (Figure 4A). The 3D structure of the protein alpha-amylase_720 was predicted using AlphaFold (Figure 4B). Homology modelling based on the recently solved structure of CBM74 from *Ruminococcus bromii* revealed the presence of a substrate analogue (a maltodecoase double helix) aligned in the CBM74 module binding pocket, positioned near the active site of the alpha-amylase. Alpha-amylase_720 shares 81.5% amino acid identity and a very similar domain structure to the previously characterised resistant-starch amylase FMB-RSD3 from *Bifidobacterium choerinum* [62]. Alpha-amylase_720 possesses a TAT signal peptide, suggesting extracellular export, and S-layer homology (SLH) domains, which anchor proteins to the bacterial G(+) cell wall. Full functional predictions are detailed in Supplementary Table S7. A BLAST search using the NCBI database, revealed that homologous proteins to alpha-amylase_720 are almost exclusively found among members of the *Bifidobacterium* genus, with *B. pseudolongum* the most common host for this protein, indicating it is conserved within this species (Supplementary Table S8). This CBM74 module is prevalent across the *Bifidobacterium* genus, and several homologs of alpha-amylase_720 in human isolates were identified including 4 strains of *B. pseudocatenulatum* and 7 *B. adolescentis* strains, each exhibiting protein sequence identities above 64.5% (Supplementary Table S8).

**Figure 4.**
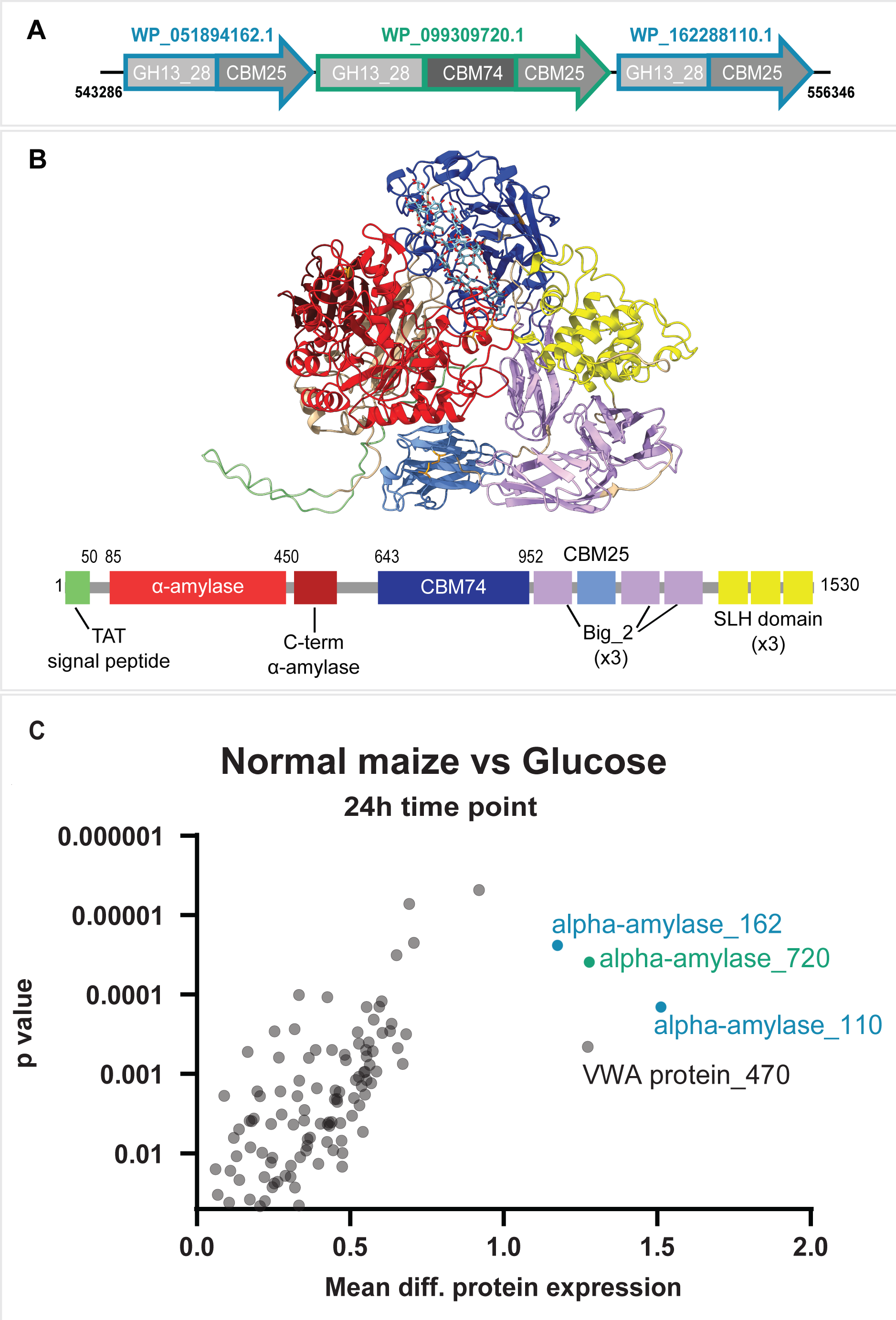
Protein and proteome analysis of *B. globosum* 45. A) The gene cluster present in the *B. globosum* 45 genome containing 3 Ig-like fold containing proteins possess a-amylase (GH13) gene CAZyme predictions and occur at the same locus in the genome. Each a-amylase is also predicted to have at least one associated starch-binding carbohydrate module (CBM). B) The central protein (WP_099309720.1) is a large multi-modular protein; its 3D structure of is shown as predicted using AlphaFold. It is predicted to contain a-amylase, CBM74, CBM25, and an S-layer homology domain. C) Differential protein production were analysed via t-test to measure significant differentially abundant proteins in the presence of 1% w/v normal maize starch vs 1% w/v glucose after 24h culture. Bacterial cultures were carried out in triplicate. The proteins highlighted correspond to the following genes: WP_051894162.1 = alpha-amylase_162; WP_099309720.1 = alpha-amylase_720; WP_162288110.1 = alpha-amylase_110.

### Proteomic analysis of a bifidobacterial isolate with an enhanced RS degrading phenotype

To assess the role of proteins in the degradation of resistant starch, we performed proteomic analysis on *B. globosum* 45 cultured in the presence of NMS compared to glucose. Of 1570 predicted proteins in the genome, 1081 proteins were detected by LC-ESI-MS/MS, and 729 differentially expressed proteins between the two conditions were detected. A total of 619 proteins were significantly upregulated in glucose condition, whilst 110 were significantly upregulated in NMS. The most differentially abundant proteins in the starch condition detected were the three proteins belonging to the gene cluster of Ig-like fold containing proteins predicted to have a-amylase activity and starch-binding modules: alpha-amylase_162, alpha-amylase_720, and alpha-amylase_110 (Figure 4C). Additionally, a vWA-domain containing protein WP_026643470.1, here called vWA protein_470, was also highly upregulated; which are commonly associated with multi-protein complexes and play roles in protein folding [63] (Figure 4C). Both alpha-amylase_162 and alpha-amylase_720 were shown to be expressed in the glucose condition, but their expression levels were significantly upregulated in the presence of starch (Supplementary Figure S1). In contrast, alpha-amylase_110 and vWA protein_470, were expressed at much lower levels in the starch condition, and were barely detectable in the glucose condition (Supplementary Figure S1). This suggests that these proteins, particularly those from the a-amylase cluster, play a key role in RS degradation, with their activity specifically induced in response to RS presence.

### *Bifidobacterium* amylase genes are differentially expressed dependent on starch structure

The total intracellular RNA of *B. globosum* 45 was measured in the presence of NMS, high amylose starch HylonVII, glucose, or in the absence of a substrate. Two time points were selected (12h, 24h) to capture different phases of bacterial growth on these substrates.

Principal component analysis showed that the largest source of observed variance in the transcriptome could be attributed to the presence of either NMS or HylonVII compared to either glucose, no substrate, or prior to introduction of starch (the inoculum culture) (Figure 5A). There was minor separation observed between NMS and HylonVII, which was also observed in a Pearson’s correlation heat map (Figure 5B).

**Figure 5.**
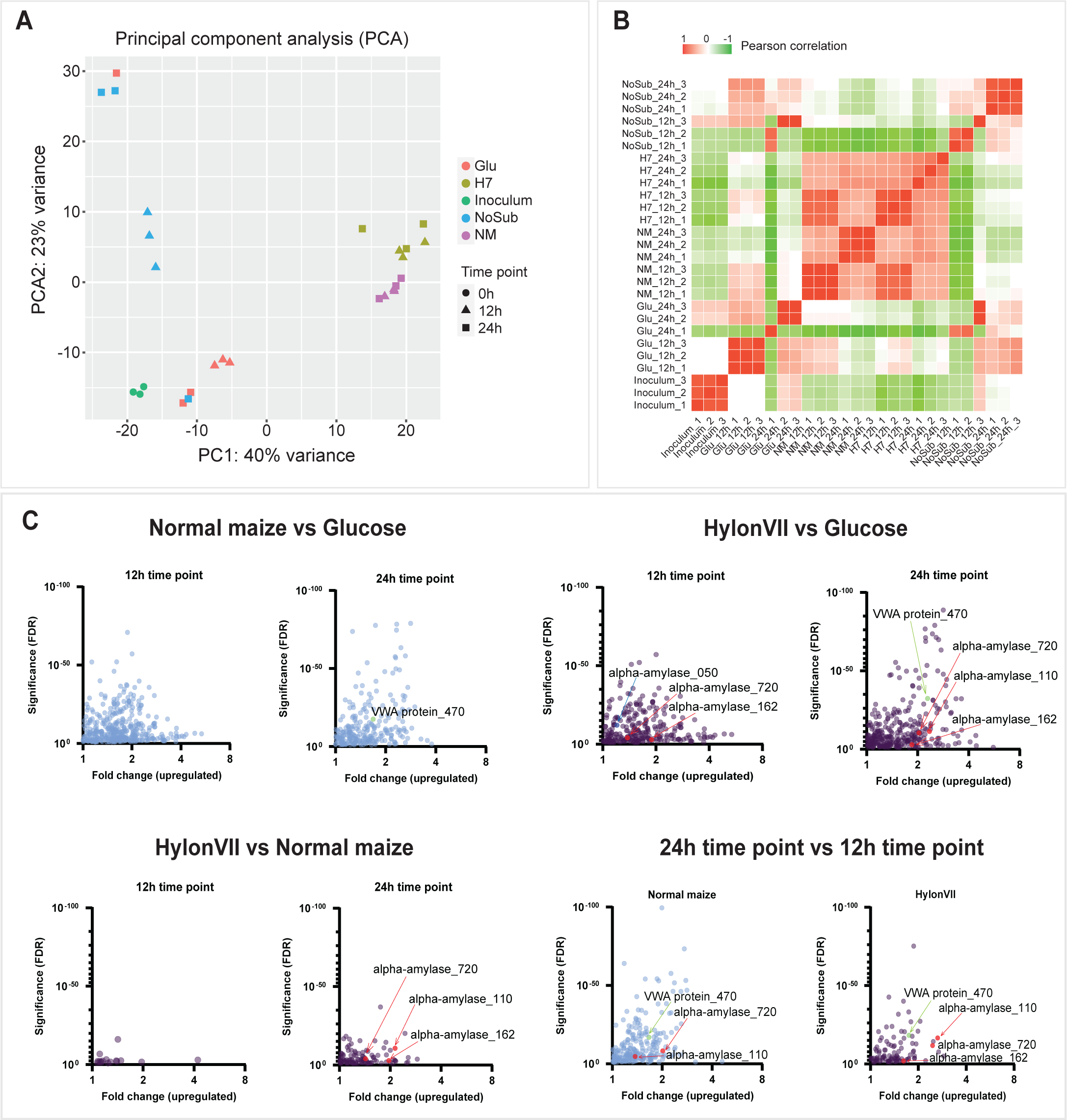
Transcriptome analysis of *B. globosum* 45 in the presence of different starch types. A) Principle component analysis of RNA expression between substrates of *B. globosum* 45. Plot displays variance of the transcriptome of strains under different conditions and time points. Substrates tested were in the presence of glucose (Glu), normal maize (NM) starch, high amylose HylonVII maize (H7) starch, and no substrate (NoSub). Time points displayed demonstrate when RNA was extracted: 0h (inoculum culture of the experiment), 12h, and 24h. B) Pearson correlation coefficient matrix describes linear correlation between transcriptome culture conditions, demonstrating differences and similarities between RNA expression in *B. globosum* 45. C) Differential genes expression volcano plots display the upregulation of specific genes, using DESeq2 to quantify fold changes in expression and significance value (false discovery rate (FDR)). Genes significantly upregulated were displayed comparing the following culture conditions: normal maize vs glucose, HylonVII vs glucose, HylonVII vs normal maize, and 24h vs 12h time point. Specific genes significantly upregulated such as vWA-domain containing gene and three clustered novel a-amylase genes are highlighted by colour.

Further differential abundance analysis using DeSeq2 was performed to quantify gene transcript levels between conditions (Figure 5C). Notably, the three putative a-amylase genes ‘Ig-like domain containing proteins’ (alpha-amylase_050, alpha-amylase_162, alpha-amylase_720), were not significantly upregulated in the NMS condition compared to glucose at either time point. However, in the presence of HylonVII during mid-exponential growth (12 h), alpha-amylase_720 was significantly upregulated (fold change = 1.38, FDR = 7.83E-05), along with another alpha-amylase at a different locus, WP_051592050.1, here called alpha-amylase_050 (fold change = 1.20, FDR = 6.77E-13). At 24 h, all three a-amylase genes in the cluster, along with the the vWA protein_470 (fold change = 2.29, FDR = 3.81E-33) were significantly upregulated in line with proteomic analysis. The a-amylase proteins and vWa protein genes are not co-located in the genome.

The differential expression of these putatively starch-degrading a-amylase genes at 12 h in HylonVII suggests that the resistant properties of this starch induced a stronger transcriptional response compared to the more accessible NMS. When comparing gene expression directly between HylonVII and NMS at each time point, significant differences were observed at 24h, particularly for the three a-amylase genes in the gene cluster.

Additionally, significant differential expression of the alpha-amylase and vWA protein gene transcripts were observed in both conditions NMS and HylonVII between the two time points (12 h *vs* 24 h) (Figure 4C). The discovery of a gene encoding a cell-wall-anchored, multi-modular a-amylase protein, flanked by CBM74 domains, and co-transcribed with two other putatively starch-binding amylase-encoding genes, represents the first such gene cluster identified in *Bifidobacterium*.

## Discussion

The effect of dietary components, age, and strain characteristics within the gut microbiota are highly relevant during infancy, a critical period of microbiome maturation [21, 64, 65]. As the infant digestive system develops, its ability to digest dietary components becomes more efficient, but is influenced by individual variation [66]. The introduction of solid foods during weaning, especially starchy substrates, stimulates the expansion of certain key taxa, like specific *Bifidobacterium* strains, that promotes further microbiota development [21]. The ability of *Bifidobacterium* species and strains to degrade starch suggests it plays a key role in microbiome transition during this pivotal developmental window, but more detailed mechanistic insights are needed to fully understand this process.

Previous research has shown that *Bifidobacterium* strains in breastfed infants engage in cross-feeding interactions, where HMO-degrading strains support the growth of non-degrading strains [29]. Our present work extends this concept, by using the same culture collection of *B. breve* and *B. pseudocatenulatum* strains, suggesting that weaning-age *Bifidobacterium* strains not only metabolise HMOs but also possess the capacity to degrade starch substrates. These starch-degrading strains were able to generate maltose and acetate as metabolic by-products, while the assessed strains that had been isolated prior to weaning could not. These findings highlight a potential bridging role for *Bifidobacterium* strains during the dietary transition from breast milk to solid foods, where they adapt to new substrates while maintaining cross-feeding interactions. This suggests that *Bifidobacterium* species may function as metabolic generalists during weaning, facilitating microbiome stability.

The diverse CAZyme profiles observed in *B. pseudocatenulatum* LH strains had the highest number of CAZymes from a variety of families such as the starch related GH13, CBM48, CBM25, and CBM74, as well as family GH43 which includes enzymes acting on a broad spectrum of substrates, providing further evidence of bacterial generalism [67]. This suggests that the *B. pseudocatenulatum* LH strains have a large variety of enzymes which would make them well suited for solid food substrates normally introduced during weaning in addition to HMOs. *B. pseudocatenulatum* as a species has also been demonstrated to have generalist phenotypes [68]. The authors identified a gene cluster in *B. pseudocatenulatum* strains which were shown to encode amylases which were not active on maltose [68]. In the *B. pseudocatenulatum* LH strains tested here, maltose accumulated in the media.

Starch structure significantly influences growth and metabolite output of saccharolytic bacteria, which is known to impact microbiome ecology [69, 70]. Indeed, the *Bifidobacterium* strains tested in this study when grown with RS such as HylonVII induced slower fermentation and prolonged acetate production. This could have important implications for modulating the microbiome ecosystem, as prolonged acetate release may enhance competitive exclusion of pathogenic bacteria and support beneficial microbial populations [11]. Our findings highlight the importance of considering starch structure in both microbiome research and the development of functional foods, particularly during the weaning window.

Two genomically closely related but phenotypically distinct isolates, *B. pseudolongum* 44 and *B. globosum* 45, were investigated using a multi-omics approach to uncover novel mechanisms of interaction between *Bifidobacterium* and starch. Both isolates are predicted to encode a number of amylase-like genes, including extracellular starch-binding amylases and intracellular amylases that target glycosidic bonds after oligosaccharides are imported into the cell. Of particular interest is the CBM74 module, a relatively recently discovered RS binding module which commonly associates with multi-domain a-amylases. These modules often work synergistically with flanking CMB25 or CBM26 domains, enabling effective docking to RS granules [27, 60, 71]. The protein structure of CBM74 has only recently been solved, revealing a unique topological arrangement, potentially enhancing its binding capacity. For example, in *R. bromii*, CBM26 was shown to bind short malto-oligosaccharides, while the CBM74 module binds to single and double helical starch and is capable of tightly binding to crystalline starches [31, 32].

The CBM74 family is found almost exclusively in *Bifidobacterium*; of the 71 gut microbes that encode CBM74 family members, 50 belong to the genus *Bifidobacterium* [72]. Despite this, relatively few investigations have explored bifidobacteria which contain CBM74 domain-containing amylases and their ability to enhance RS binding or degradation. One recent study utilised a recombinantly expressed CBM74-amylase protein from *B. adolescentis* P2P3, demonstrating the protein’s ability to degrade high-amylose corn starch granules [73]. Hybrid metagenome assemblies have also confirmed a higher abundance of CBM74 sequences in response to RS being present in an *in vitro* fermentation model [70]. In our study, dbCAN annotations revealed several ‘Ig-like domain containing protein’ genes in the stronger RS-degrading isolate *B. globosum* 45. These proteins had not previously been demonstrated in *Bifidobacterium* as being related to start binding or degradation. Ig-like domain proteins have been reported in other important gut species such as *Roseburia inulinovorans* and *Eubacterium rectal*e, where they are important for substrate binding [27]. These bacteria produce a cell-wall anchored GH13 amylase with at least one CBM26 domain and Ig-like domains, which may function as unidentified starch-specific CBMs or play a structural role in degradation. In the prolific carbohydrate degrader *Bacteroides thetaiotaomicron*, elements of its large starch uptake system contain several starch-binding domains, with a canonical Ig-like/β-sandwich fold attached to starch-active enzymes, further suggesting that this mechanism may be more widespread across gut bacteria. [27].

We found that proteins encoded in the novel gene cluster in *B. globosum* 45 (three Ig-like fold domain containing proteins) were also expressed in the glucose condition. This finding supports the growth pattern in which bacteria exhibit two exponential phases, suggesting that presence of RS triggered the transcription and translation of these proteins.

Furthermore, *B. globosum* 45’s metabolism of HylonVII starch resulted in a higher degree of transcription of the novel a-amylases, indicating that their expression is influenced by time in culture and starch structure. Importantly, homologs of alpha-amylase_720 are found in human isolates of *B. pseudocatenulatum* and 7 *B. adolescentis* making this finding relevant to human hosts. This highlights a unique mechanism by which *Bifidobacterium* can degrade resistant starches, potentially contributing to their ecological role in the gut microbiota.

The identification of vWA-domain containing proteins as part of the starch degradation machinery raises intriguing questions about their role in *Bifidobacterium*. These proteins are often involved in protein-protein interactions and may facilitate the assembly of multi-enzyme complexes for efficient starch degradation [74–76]. Additionally, their involvement in biofilm formation and host adhesion suggests they could contribute to long-term colonization and microbiome stability [77]. However, further research is needed to elucidate the specific functions of vWA proteins in both starch metabolism and host interaction.

This study provides new insights into how *Bifidobacterium* species interact with dietary starch during early life, identifying key molecular players involved in RS degradation. The early life microbiota may possess *Bifidobacterium* with multi-tasking metabolic capabilities, including starch metabolism which is a key weaning food. The discovery of novel amylase genes and the importance of CBM74 in starch binding extends our understanding of *Bifidobacterium*’s role in shaping the gut microbiome during weaning. These findings have implications for the development of bifidogenic functional foods, including incorporation of RS, synbiotics, and strategies for promoting gut health throughout life.

There are limitations in this study that should be considered. In favour of in-depth research into molecular mechanisms, the study focused on a small set of human and animal bacterial species and strains; the methodology and study design used could be tested on a wider set of isolates, in particular more pre- and post-weaning isolate pairs from the same infant. The benefits of RS degradation and acetate production by these strains was not explored *in vivo* or using cell culture methods as this was considered to be outside the scope of this work. Finally, the role of upregulated vWA protein in starch degradation by bifidobacteria was not resolved and could be the subject of future research.

## Acknowledgements

We acknowledge and thank Ingredion Inc. for gifting of HylonVII starch. The authors would like to acknowledge several people who were instrumental to completion of this work. Thank you to Adam Gicgier for his contributions to the visualisation and module prediction of the amylase gene. We would also like to thank David Baker, Rhiannon Evans, and Sumeet Tiwari of the Quadram Institute Bioscience Core Sequencing and Bioinformatics departments for their work and continuing support. This research was also supported in part by the NBI Research Computing through the provision of a high-performance computing cluster. For proteomics work completed, the authors thank Verena Breitner for excellent technical assistance. Additionally, the Exploris 480 mass spectrometer was funded in part by the German Research Foundation (INST 95/1435-1 FUGG). The authors would also like to thank Chen Meng for use of the data visualisation platform omicsViewer (DOI: 10.1101/2022.03.10.483845). Finally, we thank Andrew Jermy for his editorial help.

## Author Contributions

MEM, FJW, and LJH conceived the research with support from HCH. MEM designed the experiments with inputs from FJW, LJH, and HCH. MEM performed all the experiments with contributions from TTK, MA, and HCH. AT performed read mapping of transcriptome reads to reference genomes and read count data was analysed by MEM. RK completed the whole genome assembly. TTK and YZK developed the protocols used for NMR metabolomics. MEM analysed the NMR data with inputs from FJW and TTK. MA lead the development of protocols for bacterial protein extraction and completion of LC-MS/MS experiments along with CL. MA and CL carried out the LC-MS/MS experiments and subsequent data processing. Proteomics data was analysed by MEM with inputs from MA. DVS kindly contributed the *B. breve* type strain UCC2003. MEM, FJW, and LJH wrote the manuscript with inputs from HCH, MA, TTK, YZK, DVS, and AT.

## Data Availability Statement

The RNA-seq raw reads data can be found in the National Centre for Biotechnology Information (NCBI) database under BioProject ID: PRJNA1156008 (https://dataview.ncbi.nlm.nih.gov/object/PRJNA1156008?reviewer=1vmchv5baet3kmo1vuae4lvkj7 <u>[dataview.ncbi.nlm.nih.gov]</u>). The mass spectrometric raw files and the MaxQuant output files have been deposited to the ProteomeXchange Consortium via the PRIDE partner repository [78] and can be accessed using the identifier PXD056548 (https://proteomecentral.proteomexchange.org/cgi/GetDataset?ID=PXD056548, reviewer account username, password: GmiDdJNDDI2C). The genomic data underlying this article are available in NCBI repository. A full list of each bacterial strain and its accession number can be found in Supplementary Table S1. Remaining data underlying this article are available in the article and in its online supplementary material.

---

**Supplementary Figure 1.**
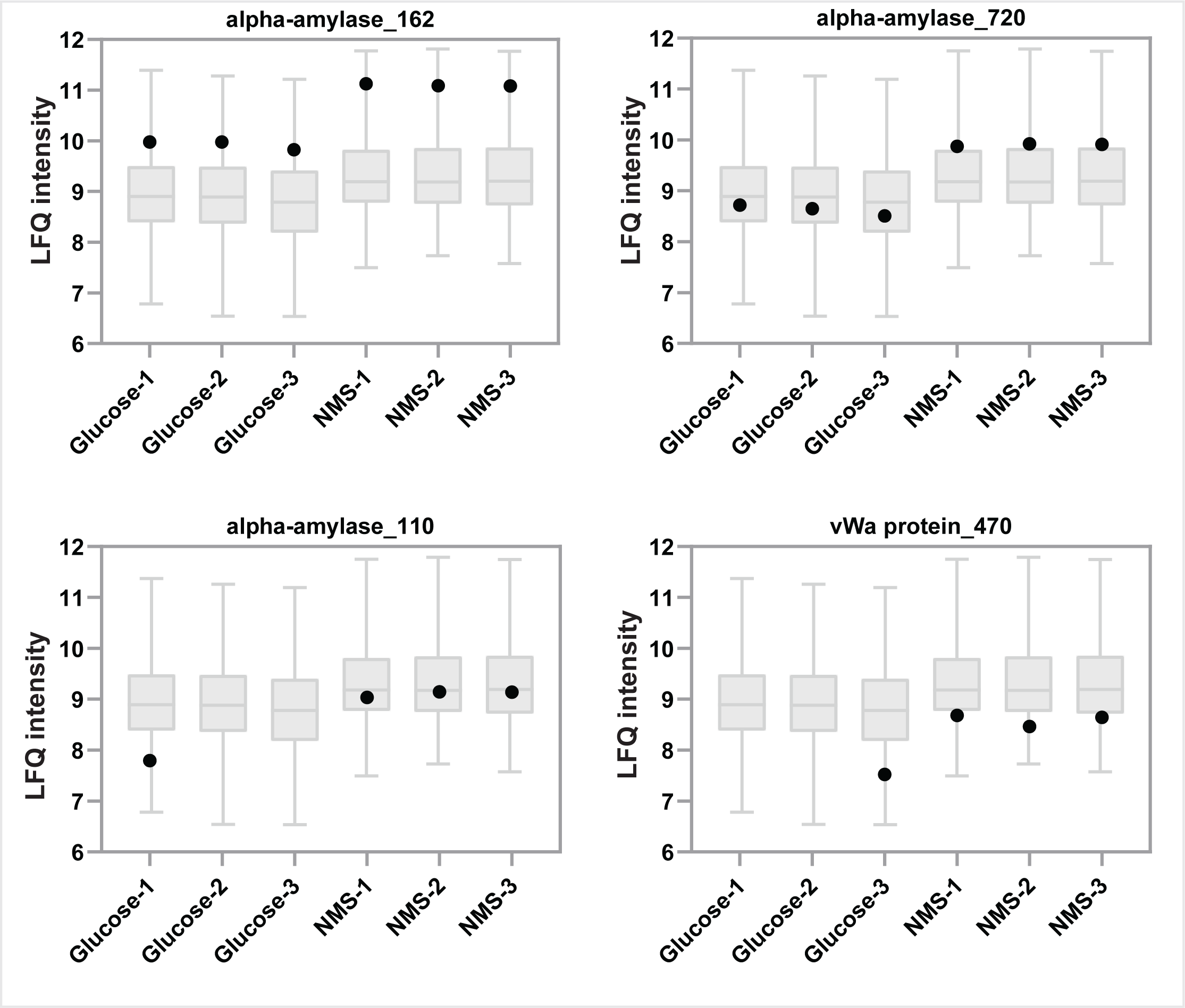
Raw LFQ intensities of individual proteins of interest. The Label-Free Quantification (LFQ) intensities of each alpha-amylase in the novel gene cluster (alpha-amylase_162, _720, and _110) and a vWa protein_470 found to be upregulated in the normal maize starch (NMS) condition compared to glucose being present in mMRS medium. Each culture condition was performed in triplicate: each replicate is displayed as a single point on each plot, with the boxplot in the background being the overall protein expression in each condition, with median and quantiles displayed.

